# Path analysis reveals combined winter climate and pollution effects on the survival of a marine top predator

**DOI:** 10.1101/2023.12.14.571445

**Authors:** Kate Layton-Matthews, Kjell E. Erikstad, Hanno Sandvik, Manuel Ballesteros, Kevin Hodges, Michael d.S. Mesquita, Tone K. Reiertsen, Nigel G. Yoccoz, Jan Ove Bustnes

## Abstract

Marine ecosystems are experiencing growing pressure from multiple threats caused by human activities, with far-reaching consequences for marine food webs. Determining the effects of multiple stressors is complex, in part, as they can affect different levels of biological organisation (behaviour, individual traits, demographic rates). Knowledge of the cumulative effects of stressors is key to predict the consequences for threatened populations’ viability under global change. Due to their position in the food chain, top predators such as seabirds are considered more sensitive to environmental changes. Climate change is affecting the prey resources available for seabirds, through bottom-up effects, while organic pollutants can bioaccumulate in food chains with the greatest impacts on top predators. However, knowledge of their combined effects on seabird population dynamics is lacking. Using a path analysis, we quantify the effects of both climate change and pollution, via an effect on body mass, on the survival of adult great black-backed gulls. Warmer ocean temperatures in gulls’ winter foraging areas in the North Sea were correlated with higher survival, potentially explained by shifts in prey availability associated with global climate change. We also found support for indirect negative effects of organochlorines, highly toxic pollutants to seabirds, on survival acting through a negative effect on body mass. The results from this path analysis highlight how, even for such long-lived species where variance in survival tends to be limited, two stressors still have had a marked influence on adult survival and illustrate the potential of such models to improve predictions of population variability under multiple stressors.

## Introduction

Natural populations are facing increasing pressure from anthropogenic stressors, which often act in concert (Jackson et al. 2021, Simmons et al. 2021). Multiple stressors can interact in complex ways, potentially combining to have negative impacts greater than the sum of their individual effects (Darling and Côté 2008). Understanding their combined effects is therefore crucial to assess the consequences of global change and to predict the future dynamics of threatened populations. Our knowledge of when and how interactions arise is limited because most models and experiments only consider the effect of single stressor, which can lead to underestimation, or overestimation, of the threats to biodiversity (Simmons et al. 2021).

Marine ecosystems are experiencing intense and growing pressure from human activities (OSPAR 2009). Overfishing, climate change, and the presence of organic pollutants have irreversibly altered the structure and stability of marine food webs (Halpern et al. 2008, Maxwell et al. 2013). Top predators are particularly sensitive to such anthropogenic impacts, as a result of their life history characteristics such as long lifespans and delayed maturation (Heithaus et al. 2008). Marine predators also tend to be bottom-up driven and therefore more vulnerable to changes in food resources (Frederiksen et al. 2006). Seabirds are top predators in most marine ecosystems. They are also considered the world’s most threatened bird group and many populations are undergoing strong declines (Croxall et al. 2012). There are numerous well documented threats to seabirds, including pollution, climate change, bycatch, and overfishing (Cury et al. 2011, Dias et al. 2019).

There is a growing evidence for the widespread, detrimental consequences of climate change for seabirds (Grémillet and Boulinier 2009, Sydeman et al. 2012). Climate change is largely affecting seabirds through bottom-up effects on prey resources (Frederiksen et al. 2006, Reiertsen et al. 2014). However, phenomena such as increased extreme weather events can also directly affect mortality rates (Guéry et al. 2019). Seabirds are considered at greater risk of climate change because of their migratory nature, which means they experience different environmental conditions at discrete sites and are therefore less able to predict changes in conditions. Particularly in Arctic regions, where global warming is occurring at a faster rate (Rantanen et al. 2022).

While pollution is considered an important threat to seabirds, our understanding of its impact on population dynamics is limited, particularly in the context of multiple stressors (but see e.g., Bustnes et al. 2015, Bårdsen and Bustnes 2022). The most damaging pollutants to wildlife, and top predators in particular, are those that persist and bioaccumulate, with particularly severe consequences higher up the food chain. For example, methylmercury, a potent neurotoxin which bioaccumulates in marine food chains, has been found at extremely high concentrations in predatory fish (Schartup et al. 2019). Persistent organic pollutants (POPs) are known to negatively affect seabirds. Individuals with higher levels of POPs can have increased parasite loads (Sagerup et al. 2000) and reduced individual condition (Bustnes et al. 2002). This, in turn, has negative implications for their reproduction (Bustnes et al. 2003), survival (Erikstad et al. 2013, Goutte et al. 2015) and population viability (Bustnes et al. 2003). Subarctic and arctic marine ecosystems currently have relatively high levels of several long transported persistent POPs (Burkow and Kallenborn 2000) and even low levels of POPs can have ecological consequences, especially under poor environmental conditions (Bustnes et al. 2015, Goutte et al. 2015).

Pollutants are released into the blood when fats are metabolised and redistributed into vital organs (Bustnes et al. 2012). Therefore, individuals’ physiological state also determines circulating levels of pollutants and thereby mediates potential negative effects on survival or reproduction (Henriksen et al. 1998, Bustnes et al. 2015). Consequently, other stressors, e.g., climate change, may exacerbate the negative impacts of pollution by affecting individuals’ condition. Conversely, contaminant exposure can also alter individual responses to environmental conditions, via regulation of stress hormones (e.g., Nordstad et al. 2012). Consequently, we need to study the combined effects of climate change and pollution on individuals’ condition and demographic rates to predict their impact on population viability. Such mechanisms may be particularly relevant for Arctic breeding seabird populations, where large fat reserves are accumulated prior to the breeding season (Henriksen et al. 1998).

Determining relationships between two main threats to seabirds (non-breeding season climate and pollution) and demographic rates, via an indirect pathway, requires an analytical approach that accommodates direct and indirect relationships among multiple variables. Path analysis is a multivariate regression technique to formalise and confront different hypothesised scenarios by correlating different variables (Shipley 1997). In this study, we adopted the framework from Gimenez et al. (2012) to integrate mark-recapture data from a long-lived seabird, great black-backed gulls (*Larus marinus*), with known individual contaminant (organochlorine) levels and climate data from their known non-breeding areas, to disentangle their effects on survival. We tested for direct effects of ocean warming and indirect effects of organochlorine contamination, via individual condition, on adult survival rates.

## Methods

### Study system and species

Great black-backed (hereon ‘GBB’) gulls are coastal species with a recorded lifespan of 10–23 years. They lay clutches of 1–3 eggs and typically begin breeding at 4–5 years old. GBB gulls are top predators and their diet consists mainly of marine invertebrates and fish, as well as other seabirds’ eggs and chicks, during the breeding season (Furness and Barrett 1985). Seabirds deposit a portion of contaminants in their eggs (Verboven et al. 2009). Therefore, gulls feeding on eggs are at even greater risk of accumulating heavy pollutant loads (Bustnes et al. 2000). The study was carried out at a seabird breeding colony, Hornøya (70° 23’ N, 31° 09’ E), in the southern Barents Sea. Data have been collected from a great black-backed (GBB) gull population breeding at Hornøya since 2001. There were an estimated 200 breeding pairs in 2001 and 113 pairs in 2018 (Reiertsen, pers. comm.), i.e., an almost 50% reduction.

### Biometric and pollutant data

Breeding adults were caught using nest traps during the incubation period (May–June) in 2001 and 2002. Once caught, birds were weighed, and the head and bill length were measured. GBB gulls were sexed using measurements of head and bill length as they are size dimorphic (see Bustnes et al. 2008a for details). Individuals’ body mass (weight) was used as a measure of individual quality.

Organochlorines (OCs) are chemical pesticides and are distributed globally with toxic, biomagnifying effects on wildlife (Jones and De Voogt 1999). Of all persistent organic pollutants, OCs are considered to have the most detrimental effects on marine top predators (Bustnes et al. 2008, Murphy et al. 2018). Although OCs are no longer in use, these compounds are persistent and are still contaminating terrestrial and aquatic environments worldwide (Buccini 2003). Levels of several OCs in the blood were measured from 158 different individuals during field campaigns in 2001–2002. Approximately 10ml of blood was drawn from the brachial vein. Levels of organochlorines were analysed at the Environmental Toxicology Laboratory at the Norwegian School of Veterinary Science (see Bustnes et al. 2008a). Samples were analysed for hexachlorobenzene (HCB), p,p’-dichlorodiphenyldichloroethylene (DDE), oxychlordane and the total polychlorinated biphenyl (PCB) congeners. Blood concentration (ng/g wet weight) was used as a measure of an individual’s relative OC levels. Other studies suggest a relatively high stability of OC blood concentrations in incubating gulls, under stable conditions, indicating that blood concentration is a reliable measurement of individuals’ relative OC burden (Bustnes et al. 2001, Bustnes et al. 2005). Data of OC concentrations had a left-skewed distribution and so were log transformed to approximate a normal distribution (Supplementary Information S1 Figure S1). Each OC variable was then normalised (mean equal to 0 and standard deviation equal to 1) within year (separately for values from 2001 and 2002) to account for differences in the timing of capture and total levels of contaminants between years. The first principal component (PC1) from a principal component analysis (PCA) was used to represent the overall contamination level of each individual (Supplementary Information S1 Figure S2). The PCA was applied to the log-transformed and scaled values of the measured compounds (HCB, DDE, summed PCBs and oxychlordane), using the ‘prcomp’ function in R (version 4.2.2, R Core Team 2023). The first axis of the PCA, PC1, explained 88% of the total variance in OC concentrations.

### Non-breeding season distribution

We estimated gulls’ core foraging areas during the non-breeding season using geolocation data from 2012–2015 (Figure 1). Ten adult birds were equipped with miniature year-round, light-based tracking devices (‘geolocators’), which were attached to the plastic leg ring. Light data from the geolocators were processed using the BASTrak software package (British Antarctic Survey, Fox 2010) and positions were calculated using a threshold method (Lisovski et al. 2020). Then, raw positions were filtered with distance, speed, loess filters and unreliable positions around the equinoxes, between 7^th^ September–19^th^ October and 24^th^ February–5^th^ April, were excluded. Utilisation distributions (10%, 30%, 50% and 70%) were estimated as kernel density distributions using the “kernelUD” function in the “adehabitatHR” package (Calenge 2006). Based on the geolocation data, individuals also forage close to the breeding colony at Hornøya during spring prior to breeding (Figure 1). After breeding, they travel to the North Sea around the east coast of the UK, and most birds remain there during the winter. During late winter, individuals migrate back to the southern Barents Sea area (Figure 1).

**Figure 1.**
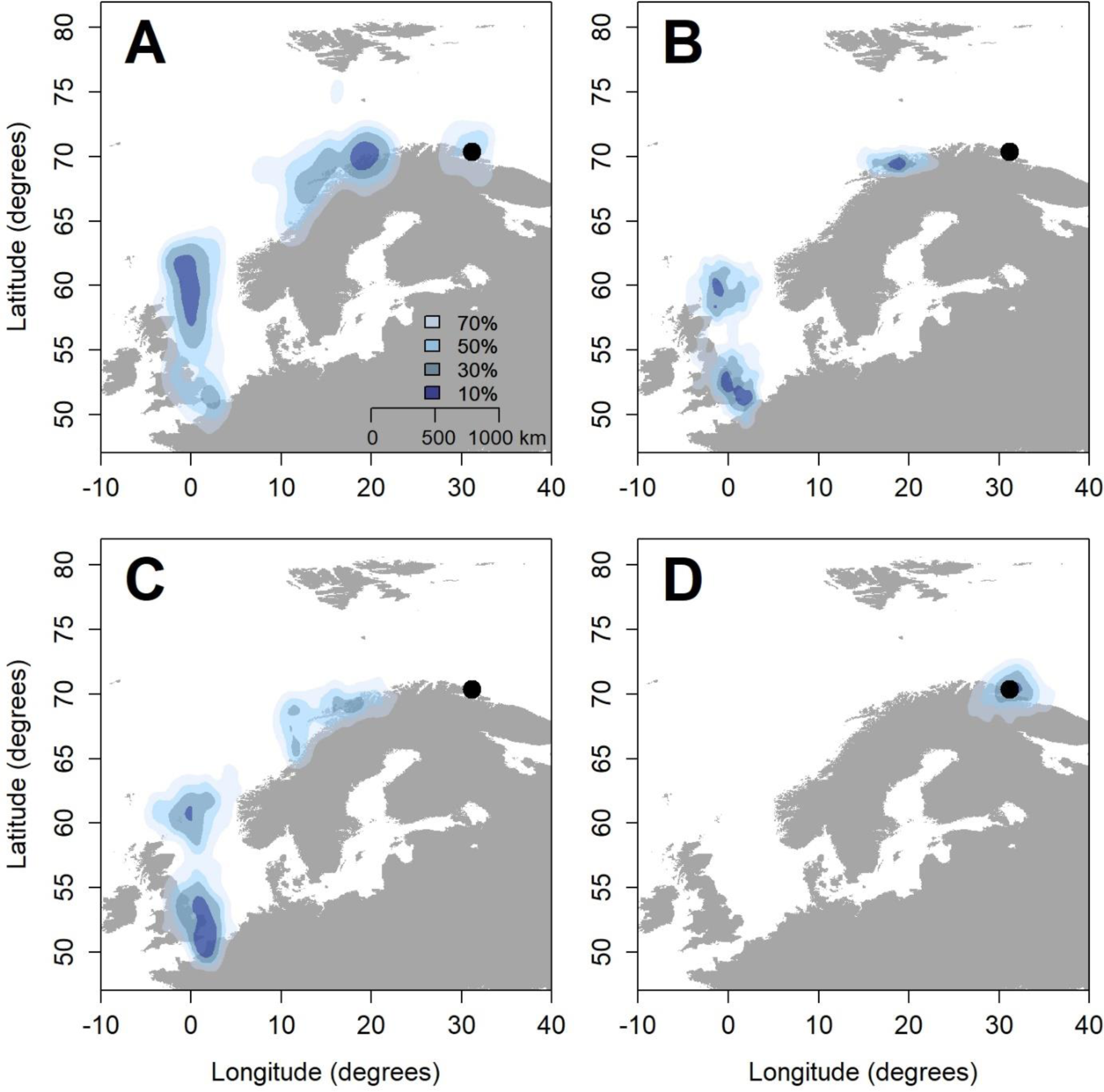
Year-round non-breeding areas of adult Great Black-Backed gulls from Hornøya based on positions from individuals with geolocators (GLS) during the years 2012-2015 for autumn (September – October, **A**), early winter (November – December, **B**), late winter (January – February, **C**) and spring (April, **D**) seasons.

### Climate data

We selected climate variables reflecting non-breeding season conditions that may affect survival rates, which were available at different spatial scales; North Atlantic Oscillation, Atlantic Water inflow, land temperature, sea surface temperature and mean sea level pressure.

North Atlantic Oscillation (NAO) and Atlantic Water inflow (AW) were available as monthly values. The NAO index reflects large-scale weather patterns and specifically cyclone activity in the North Atlantic and is widely used as a proxy for indirect effects of climate conditions on seabirds via changes in prey distributions and abundances (Stenseth et al. 2003). Annual, monthly values of NAO index are openly available from www.cpc.ncep.noaa.gov. In the Barents Sea, ocean conditions are largely determined by fluctuations in the inflow of warm and saline Atlantic water (AW; Loeng 1991, Ingvaldsen et al. 2004), again affecting seabirds through changes in prey availability (Barrett et al. 2017). Cyclones, in part, modulate influx of Atlantic water and so this can be related to NAO (Heukamp et al. 2023). Data of AW were available at a monthly scale from www.thredds.met.no/thredds/catalog/nansen-legacy-ocean/SVIM/catalog.html. NAO and AW were aggregated for two periods: winter (November–February) and spring (March–April).

Land temperature (airT), sea surface temperature (SST), mean sea level pressure (MSLP) were available as gridded data and were extracted from within seabirds’ seasonal foraging ranges (Appendix S2, Fig. S2) and were available from the European Centre for Medium-Range Weather Forecasts (ECMWF) Re-Analysis Interim Project (‘ERA-Interim’). ERA-Interim is a gridded model dataset at a resolution of 0.70° or approximately 79km based on data assimilation of meteorological station data, satellite data, among others (Dee et al., 2011; Mesquita et al., 2015). Values of MSLP, SST and airT were extracted from areas reflecting the utilisation distributions (Figure 1) for the relevant seasonal periods (Supplementary Information S2 Figure S3). Annual covariates were aggregated to four periods: Autumn (September–October, Aut), early Winter (November–December, Ewin), late Winter (January– February, Lwin) and Spring (March–April, Spr), for the years 2001–2017. Values were extracted from the North Sea region for Autumn, early Winter and late Winter (NNS/SNS, Figure S3). For spring, values were extracted from around the breeding colony at Hornøya (HRN, Figure S3). See Supplementary information S2 Figure S4 for plots of the annual time series.

### Survival model selection with climate covariates

We used individual-level capture-mark-resight (CMR) data collected between 2001 and 2017. Only mark-recapture data from individuals with known levels of OCs (N=158), i.e., caught and sampled in either 2001 or 2002, were included in the analysis. During first capture, birds were marked with a unique numbered metal ring and individually coded colour ring for future resighting, which is possible with a telescope or binoculars. Each year, visual searches were made for marked birds, predominantly at the same colony where initial capture and ring-marking took place.

The CMR analysis was based on a Cormack–Jolly–Seber (CJS) model framework (Lebreton et al. 1992). Apparent survival and re-sighting rates were estimated using MARK via the program RMark (Laake 2013) run in R (version 4.2.2, R Core Team 2023). The goodness of fit (GOF) of this model to the data was assessed using UCARE (Choquet et al. 2009). One of the GOF test components indicated a significant deviation from the model assumptions: Test 2.CT showed trap-dependence (dependence of resighting rates across years, χ^2^ = 40.86, df = 10, p < 0.001). We corrected for this by including a trap-dependence effect in the resighting model, where resighting rates of individuals the year after first capture are estimated separately from those that were not resighted the previous year. Correcting for this effect sufficiently improved the model fit (χ^2^ = 12.30, df = 10, p = 0.27).

We first obtained the best structure for a model without covariates (Supplementary Information S3 Table S1). Given the large number of climate variables, candidate models of survival were then fitted which included climate variables. Model selection was based on Akaike’s Information Criterion corrected for small sample size (AIC_C_; Burnham and Anderson 2002). A maximum of two covariates per model was set and variables with a correlation of 0.4 or higher were not included in the same model. Models with lower AIC_C_ are preferred; if models differ by less than 2 AIC_C_ units, they are regarded as equally well supported (unless they differ in the number or parameters, in which case the most parsimonious models within this set were favoured). We report the difference (ΔAIC_C_) between a the AIC_C_ of a given model and the AIC_C_ of the null model. The proportion of variation explained by a given survival model, R^2^, is given by;

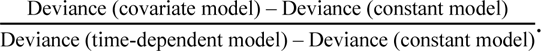

Since some of the climate covariates displayed temporal trends, models that had a lower AICc than the intercept model were tested in a de-trended version as well, to ensure effects were not spurious (see Supplementary Information S3 Table S2). In this case, covariates were linearly detrended using the ‘detrend’ function in the R package ‘*pracma*’ (Borchers 2019).

### Model selection for body mass

To determine the best model of body mass, we formulated candidate models including only an intercept term, OC level, sex (female/male) and a model with an interaction between OC level and sex. As oxychlordane is considered an especially toxic organochlorine compound (Bustnes 2006), we tested for both effects of oxychlordane levels alone and PC1 of all measured compounds. Model fits were also compared using the AIC_C_ described above.

### Path model

The path model was fitted to individual capture histories and implemented in a Bayesian framework using Markov chain Monte Carlo (MCMC) simulations to estimate posterior distributions of model parameters, following the methodology outlined in Gimenez et al. (2012). The path model estimates survival (Φ*_i,t_*) and resighting probabilities *p_i,t_*, which included a year and a trap dependence (TD) effect. The probability of resighting on the logit scale for individual *i* and year *t,* can be expressed as:

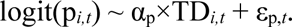

The temporal residual term, ε_p,t,_ was assumed to be normally distributed with a zero mean and variance equal to σ^2^. We developed a CMR model for GBB gulls, with a direct effect of climate conditions (time varying) and indirect effects of OC contaminant level (individually varying) on adult survival, with body mass (individually varying) as a mediating variable. Body mass was normalised so that the overall mean was zero and standard deviation of the data was 1. Total contaminant level (PC1) was fitted as an individual-level covariate (OC*_i_*). The probability of survival on the logit scale for individual *i* and year *t,* was expressed as a function of body mass (bm*_i_*) and the climate variable(s) from the best fitting model (clim*_t_*):

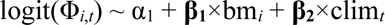

**β**s refer to regression coefficients to be estimated. Note that, through the use of a path model, body mass (bm*_i_*) was included as both a predictor variable (of adult survival) and a response variable. When fitted as a response, bm was assumed to be normally distributed with a zero mean and variance equal to σ^2^_bm_. The linear model for body mass (bm_i_) against contaminant level and sex (sex_i_) can be written as:

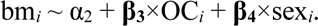

The product of **β_1_** and **β_3_** captures the indirect effect of individual level contamination on survival rates, accounting for annual variation in climate conditions and sex differences in body mass. An interaction effect between OC and sex was not included given the results from the model selection, which are shown in Table 2.

**Table 1.**
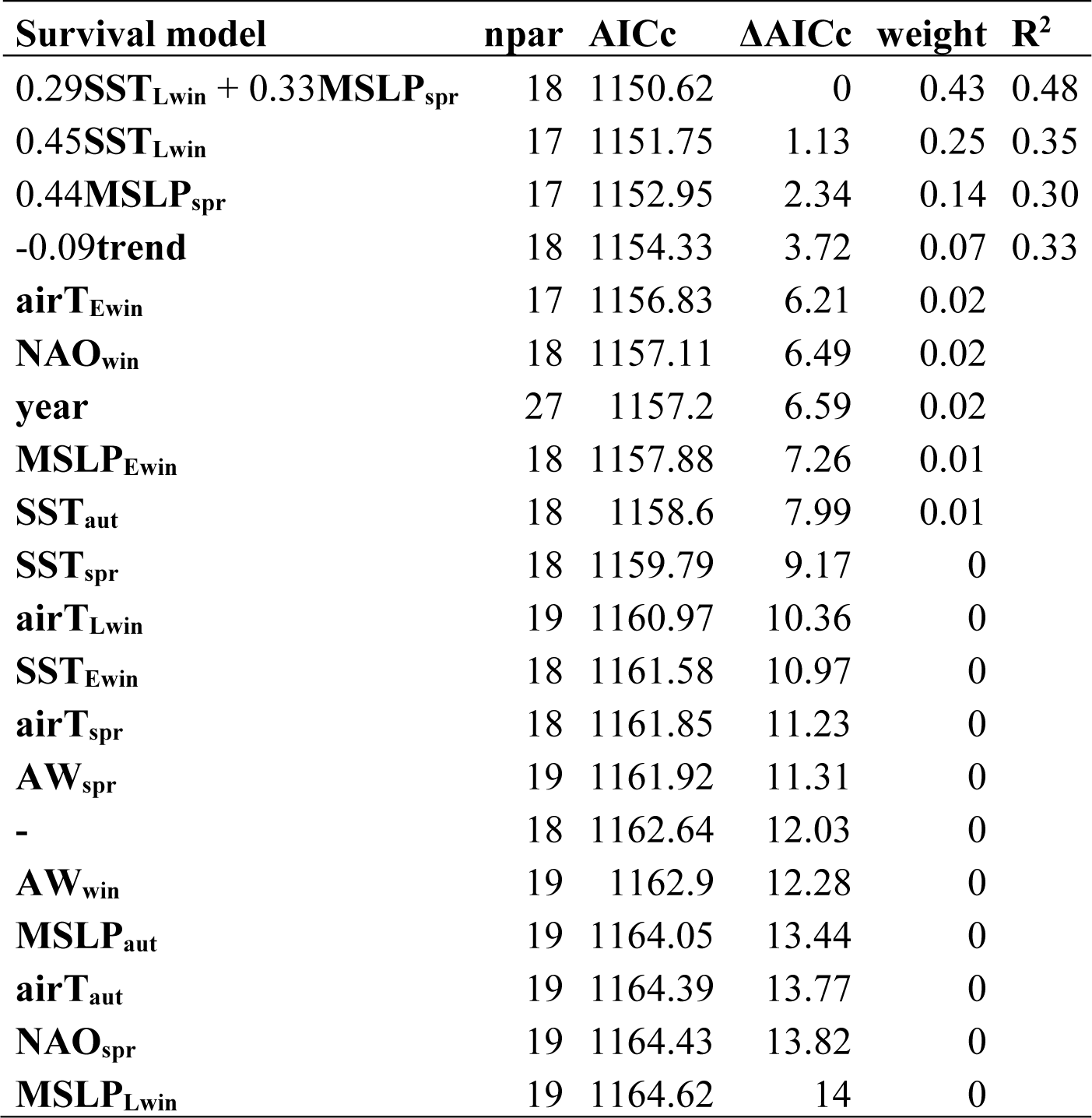
Candidate models of climate covariate effects on adult survival. For all candidate models, the recapture model included a year and trap dependence effect. Mean slope estimates are included for the covariate effects or the top models, i.e., with a lower AICc value than the trend model.

**Table 2.**
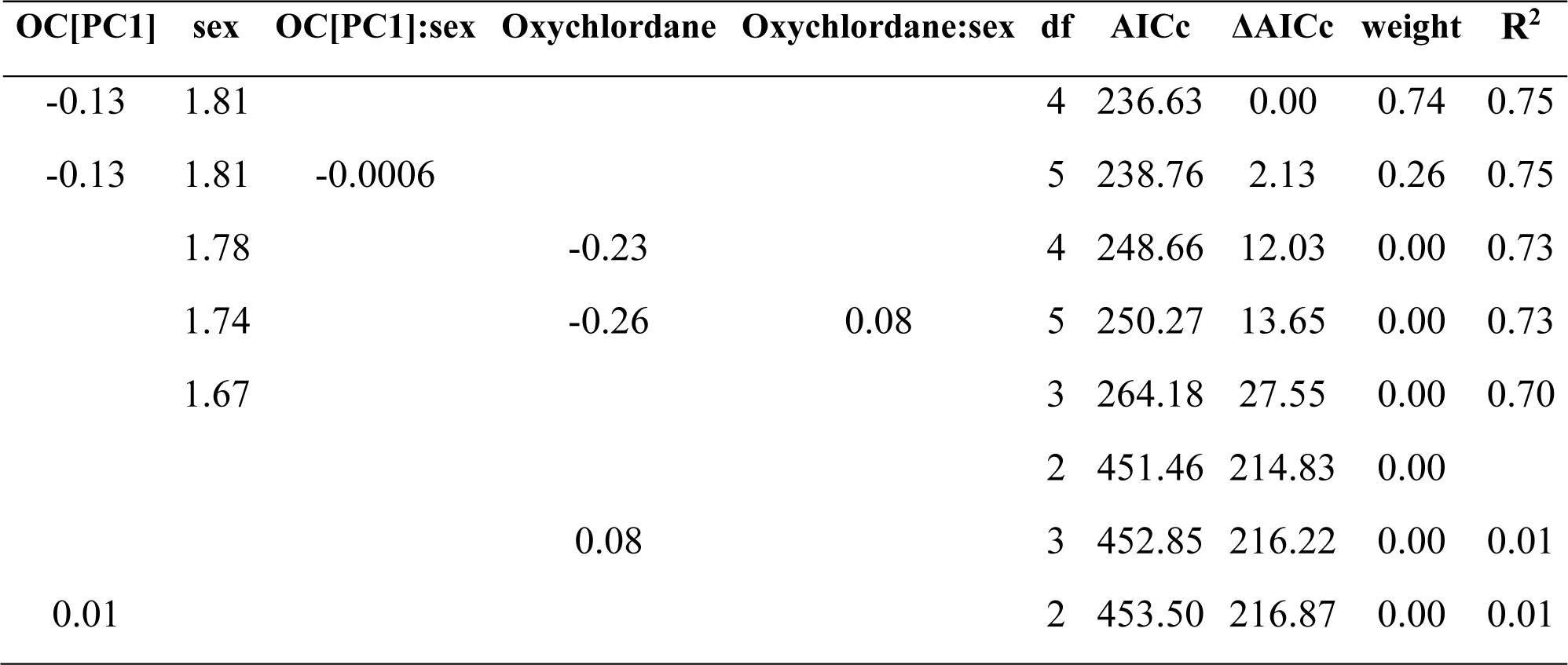
Candidate models of covariate effects on body mass. Candidate models included sex (female/male, where the slope effect is for females), and OC level and an interaction between the two terms. OC levels were represented in two ways; either as oxychlordane alone (the most toxic compound) or as the first principal component (OC[PC1]) of all compounds after log transformation.

Following Gelman et al. (2008), we used weakly informative default prior distributions for parameters to aid convergence. For the survival intercept (α_1_), a normal distribution was specified with equal to 1 and a precision of 0.37, while the mean of the normal distribution was specified as 0 for the prior for α_2_. For the regression coefficients (**β**), we specified a Cauchy prior with mean 0 and 2.5 variance, which is recommended for standardised data (Gelman et al. 2008). Normal distributions were specified with variance terms, with uniform priors with limits of 0 and 5, for the temporal random effect for resighting probability and body mass as a response. Three chains were run of length 200,000, where the first 50,000 was discarded as burn-in. Convergence was assessed by ensuring Ȓ values for each parameter were less than 1.1 (Brooks and Gelman 1998). Parameter estimates are given as means with 95% credible intervals and probabilities of posterior distributions being greater or less than zero (note posterior probabilities are rounded to two decimal places).

## Results

The best-fitting model of body mass included additive effects of PC1 and sex, but not an interaction effect between them (Table 2). Mean levels of organochlorines (OCs), measured in individuals at breeding Hornøya, were 3.611ng/g wet weight (SD = 3.06) for HCBs, 14.03ng/g (13.28) for DDEs,

76.95 ng/g (88.09) for summed PCBs and 2.18ng/g (3.13) for oxychlordane. A higher PC1 value (the first axis of the PCA where OC variables were log-transformed and scaled) was correlated with reduced body mass (-0.13 [95% confidence intervals: -0.17, -0.08]). Body mass was also higher for males than females on average (male-female contrast = 1.80 [1.64, 1.97]).

The best fitting model of survival without climate covariates, included a negative linear trend over time (Supplementary Information S3 Table S1). Based on this model, mean survival was 0.81 [0.78, 0.84] and the temporal trend was -0.09 [-0.15, -0.03] on the logit scale, where estimated apparent survival ranged from 0.86 in 2002 to 0.61 in 2017 (see Supplementary Information S3 Figure S5 for predicted survival probabilities for the trend model and recapture probabilities). The next best model of survival, which was within 2AICc units, also include an effect of sex, where females had higher survival than males (Table S1). However, the first model was the most parsimonious. The best fitting model of resighting probability included a year-specific and trap-dependence effect. The mean recapture probability was 0.45 [0.23, 0.68] and the trap dependence effect was 2.03 [1.37, 2.69] on the logit scale, indicating that birds caught the previous year were more likely to be caught the following year, i.e., ‘trap-happiness’.

The best fitting model of survival with climate covariates included a positive effect of sea surface temperature in late winter in the North Sea (SST_Lwin_, 0.29 [0.00, 0.59]) and a negative effect of mean sea-level pressure in spring around Hornøya (MSLP_spr_, 0.33 [-0.03, 0.69]), where these covariates together explained 48% of the variance. However, a model with only SST_Lwin_ (slope = 0.45 [0.14, 0.76]) was within 2AICc units and explained 35% of the annual variation in survival rates alone. This model was therefore considered the most parsimonious as it had one fewer estimated parameter. All of the described covariate models were a better fit than the baseline model of survival with a temporal trend and the intercept model (Table 1).

Based on the model selection results, a path analysis CMR model was fitted where adult survival was regressed against SST_Lwin_ and body mass (Figure 2a). Posterior distributions are shown in Figure 2b. Based on the path model, estimated mean adult survival was 0.80 [95% credible intervals: 0.77, 0.83]. Adult survival was positively correlated with body mass (Figure 3a) although 95% credible intervals overlapped zero and the posterior probability, Pr(**β_1_** > 0), was 0.81. Survival also increased with late winter SST (Pr(**β_2_** > 0) = 0.99) (Figure 3b). In the same model, body mass was also fitted as response variable with individual OC level (PC1) and sex as predictor variables. Individual body mass was lower at higher levels of organochlorines, where Pr(**β_3_** < 0) was 1.00 (Figure 2c and 3c). Body mass was also higher overall for males than females (Pr(**β_4_** > 0) was 1.00, Figure 2c and 3c).

**Figure 2.**
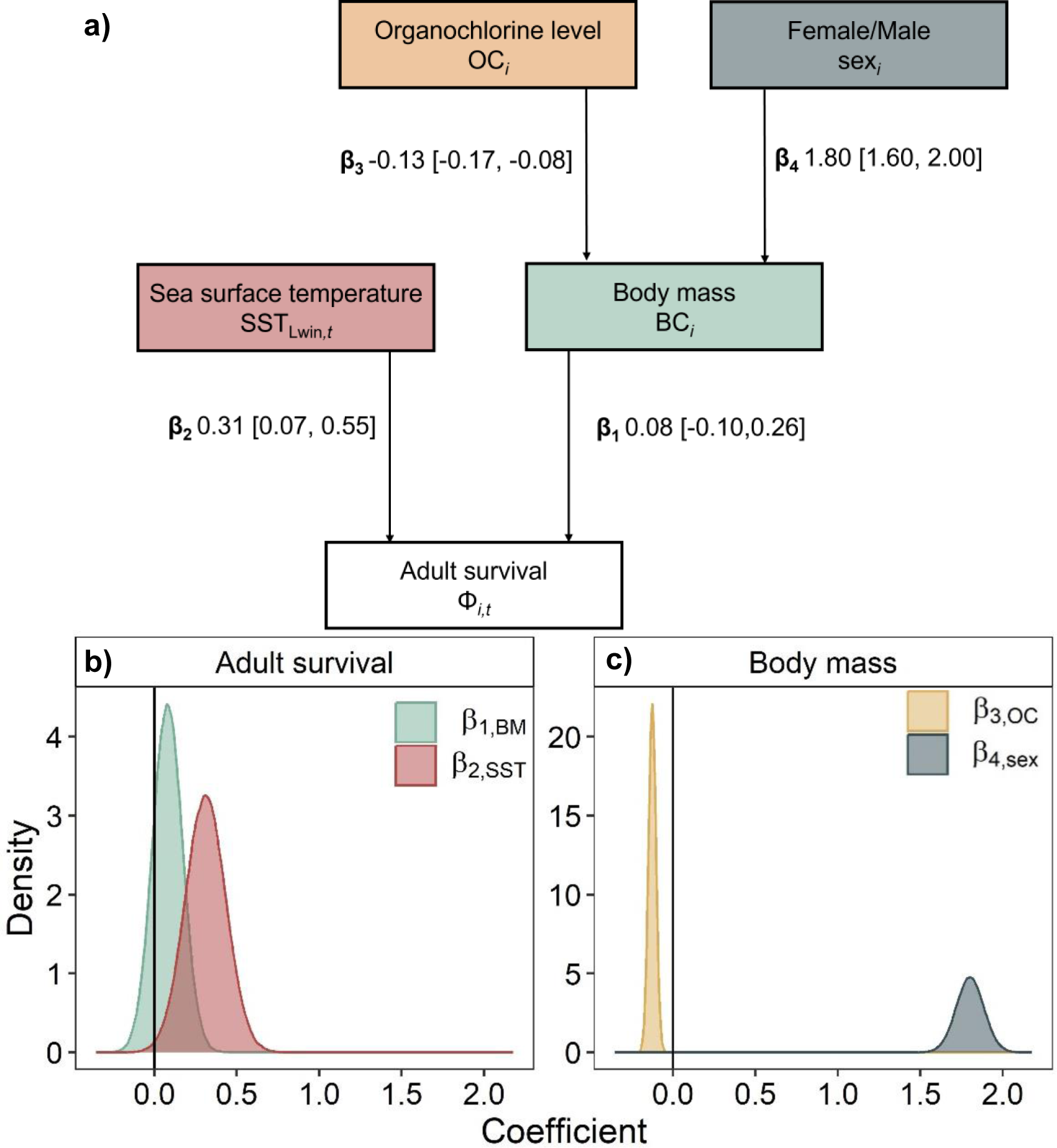
Path analysis diagram of the modelled effects of organochlorine levels and sex on body mass, and effects of SST_Lwin_ and body mass (latent variable) on adult survival. **a)** Path analysis diagram of the relationship between sea surface temperature, body mass and contaminant load, and adult survival. The indirect effect of contaminant load through body mass is captured by the product of β_1_ and β_3_. Posterior distributions of estimated regression parameters from the path model representing the effect of **b)** body mass (β_1_, green) and SST (β_2_, red) on survival rates and **c)** organochlorine levels (β_3_, yellow) and sex (slope for males = β_4_, grey) on body mass.

**Figure 3.**
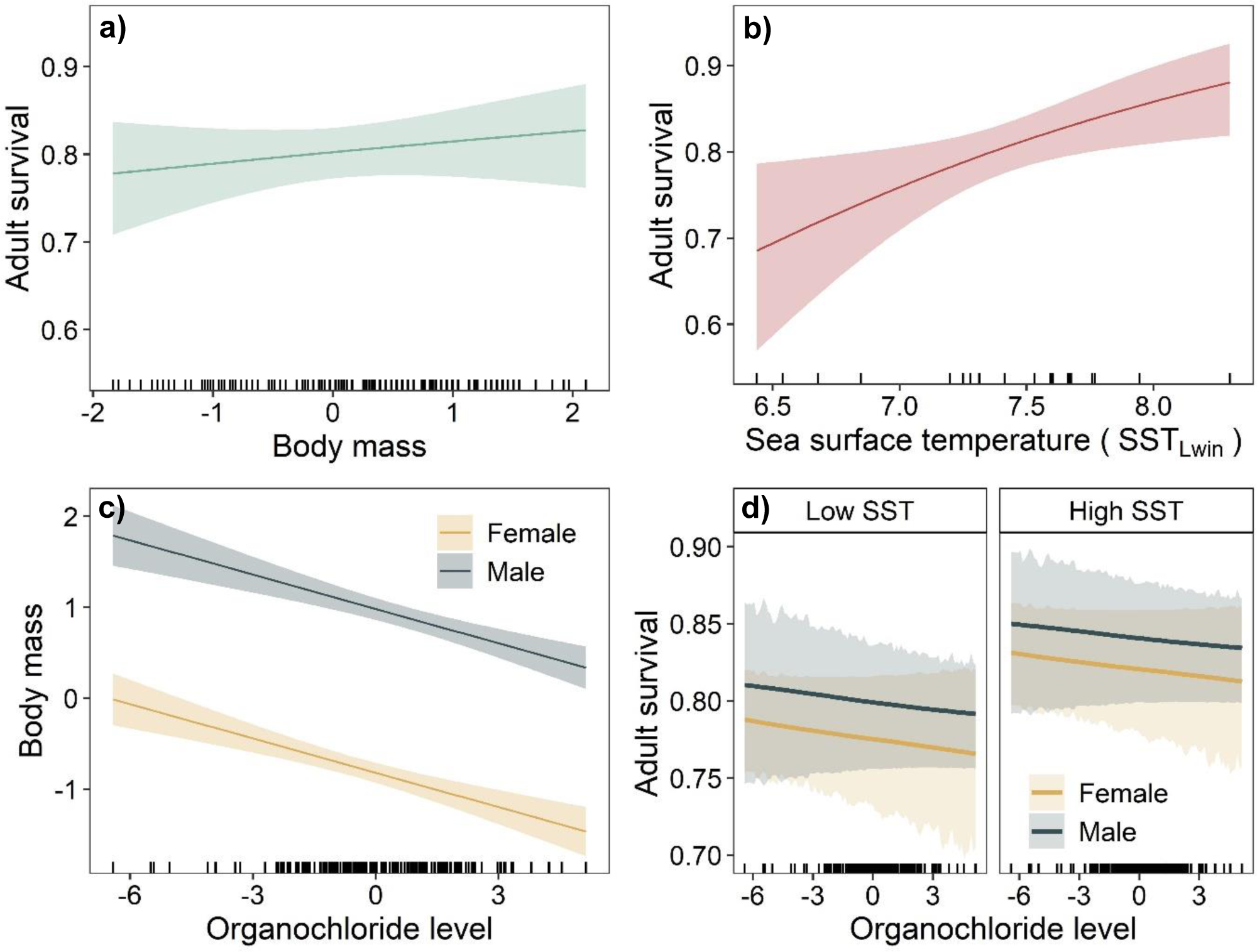
Model predictions (mean and 95% confidence intervals) for adult survival as a function of **a)** body mass and **b)** late winter sea surface temperature in the North Sea (SST_Lwin_) and **c)** scaled body mass as a function of OC level for females (yellow) and males (grey). **d)** Predictions of the indirect effect of organochlorine levels on adult survival, via a negative effect of OC on individual body mass (β_1_ × β_3_). In **d)** confidence intervals are not smooth lines as they are the product of two posterior distributions, however a smoothing function was used for the mean slope for visualisation purposes. Survival was prediction under low (25^th^ quantile) and high (75^th^ quantile) values of observed SSTs. Scaled values are shown for all variables except SST_Lwin_ in **b)**.

The product of the regression coefficients **β_1_** and **β_3_**, reflecting the indirect effect of OC levels on survival via body mass, was -0.01 [-0.04, 0.01]. An increase of one standard deviation (SD) in PC1 (1 SD = 1.87), which reflects variation in OC levels, would mean an increase in blood concentrations of contaminants (averaged across measure components HCBs, DDEs, PCBs and oxychlordane) from the overall mean 24.19ng/g wet weight to 78.31ng/g. Given the effect size of **β_1_** and **β_3_**, this would reduce survival by 0.003, i.e., a 0.37% decline. Consequently, this indicates a small, negative effect of OC levels on adult survival and also that survival was higher under favourable conditions (high SST), acknowledging parameter uncertainties (Figure 3d).

## Discussion

Anthropogenic pressure on the world’s oceans is increasing. Despite this, quantifying and predicting effects of stressors on wildlife populations in isolation, rather than their cumulative effects, remains the norm. The results from this path analysis indicate how pollution and climate affected the body mass and survival of a marine top predator, where both survival of breeding adults and numbers of breeding pairs have been in decline. While warmer ocean temperatures in the North Sea appeared to increase gull survival rates, pollution negatively, albeit weakly, affected survival, mediated by body mass.

Adult survival is a key process driving population trends in long-lived species (Sæther and Bakke 2000) and appears to have been under strong decline in this population. Therefore, this decline in adult survival probability largely explains the decline in population sizes also observed over the study period. However, given that we modelled apparent survival rates, we cannot confirm whether declines are due to true mortality or permanent emigration. Path models can be used to determine the impacts of growing pressures on marine ecosystems, through different biological pathways, and to what extent they explain declines in demographic rates and population abundances. Demographic consequences of anthropogenic stressors can be direct (e.g., mortality caused by extreme weather or bycatch by fishing vessels) or indirect, reducing an individual’s condition and thereby their future survival or reproductive potential. For instance, in Cassin’s auklets (*P. aleuticus*), availability and quality of food resources affected breeding adults’ body mass in spring and thereby their subsequent fledging success (Johns et al. 2018). Persistent organic pollutants can affect individuals’ condition, e.g., increasing levels of stress hormones (Nordstad et al. 2012) and the risk of disease (Sagerup et al. 2000, Sonne et al. 2020), and thereby reducing individuals’ condition (Bustnes et al. 2002, 2003, 2008b, Nordstad et al. 2012, Bustnes et al. 2015). Here, higher organochlorine levels were negatively correlated with individuals’ body mass during the breeding season. This is supported by studies from other gull populations, where organochlorines have been shown to reduce individual condition (Bustnes et al. 2003, Helberg et al. 2005), as well as delay laying dates and reduce clutch sizes (Helberg et al. 2005). In other colonies, especially in the high Arctic, far higher levels of organochlorines have been recorded (Steffen et al. 2006). Despite relatively low levels of pollutants at our study site in the southern Barents Sea, there was still a strong negative association between contamination and body mass (a proxy for individual quality). It is important to note that our study was an observational analysis and so we cannot infer causal links between variables. Although we modelled body mass as a function of contamination, the causal link may be the reverse: individuals experiencing mass loss due to high stress from poor environmental conditions, are known to release contaminants into the bloodstream as they burn fat reserves (discussed above). However, we do not have annual measurements of body mass, rather one measurement per individual. This means that, in this case, body mass may reflect individual quality rather than within- or between-season variation in condition. Either way, it is plausible that individuals of low quality or condition – caused by, or else evidenced by, high contamination – are more susceptible to adverse climatic conditions than individuals of higher quality or condition, leading to a cumulative effect of different types of pressures on survival.

Individual quality (body mass), measured during the breeding season, was positively correlated with higher survival. Despite having only one mass measure per individual, this path analysis provides quantitative evidence of a pathway from individual organochlorine levels to survival. In long-lived species, annual fluctuations in reproductive rates and survival of younger age classes in response to environmental conditions are often greater than in adult survival. In line with this, several studies have found evidence of negative effects on OC levels on seabird breeding success, even at relatively low contamination levels (Helberg et al. 2005, Bustnes et al. 2008a, 2008b, 2015) while evidence of pollutant effects on survival is more limited. At high Arctic Bjørnøya, organochlorine compounds in glaucous gulls have been recorded at far higher levels than those measured at Hornøya (concentrations measure at Bjørnøya in ng/g HCB = 11.3, oxychlordane = 16.7, DDE = 61.2 and PCBs = 288.9, Bustnes et al. 2003). In this study, organochlorine contamination had a stronger negative effect on survival rates than in our study (Bustnes et al. 2003, Erikstad et al. 2013). Despite the small effect on survival found in this study, likely at least in part due to the lower levels of contaminants and to having only one sampling occasion per individual, even a small change in survival has the potential to affect population viability given its importance in driving population trends, particularly when considered in the context of other stressors associated with human activities.

Identifying the environmental factors that drive population dynamics, and during which periods of the annual cycle for migratory species, is imperative, particularly for threatened populations. This population of GBB gulls, which breed in northern Norway is in rapid decline. A large proportion of this population migrates to the North Sea during the autumn and remains there until spring. The North Sea represents an important area for many seabirds and other migratory birds (Dunnet et al. 1990). Adult gulls’ survival was improved under warmer ocean temperatures during late winter in the North Sea. The Northeastern Atlantic has been strongly affected by climate change in recent decades, with associated shifts in the abundance and distribution of many species across the marine food web (Lenoir et al. 2011, Barton et al. 2016). Distributional shifts associated with climate change have been recorded at lower trophic levels which have, in turn, affected the abundance of prey for seabirds (Frederiksen et al. 2013). The consequences appear greatest for more specialised, piscivorous species (Sydeman et al. 2021). In contrast, larger gull species are generalists in their prey selection (Harris 1965, Kubetzki and Garthe 2003) and can feed on terrestrial food sources (Gyimesi et al. 2016) as well as the eggs and young of other seabirds (Harris 1965). Studies of GBB gulls indicate that their diet predominantly consists of fish and marine invertebrates, crabs in particular (Rome and Ellis 2004, Steenweg et al. 2011). Swimming crabs (such as *Liocarcinus* spp.) were also shown to be a major dietary component for lesser black-backed gulls in the south-east North Sea (Schwemmer and Garthe 2005). Warming ocean temperatures in the North Sea have been associated with increased abundances of swimming crabs and subsequent increases in the breeding success of lesser black-backed gulls (Luczak et al. 2012, Schwemmer et al. 2013). Although anecdotal, such changes in the coastal feeding ecology provide a feasible mechanism for the positive relationship between annual fluctuations in GBB gull survival and winter ocean temperatures in the North Sea.

The best-fitting, but most parsimonious, model of apparent survival also included a positive effect of mean sea level pressure in spring, around the breeding colony in the southern Barents Sea, higher MSLP reflects a greater frequency of higher-pressure systems in the area. High sea level pressure creates clear skies and blocks the passage of storms, while low pressure systems reflect stormier weather. Stormy conditions may increase energetic costs associated with foraging behaviours and can even prevent feeding for extend periods, potentially resulting in starvation (Guéry et al. 2019).

The North Sea is projected to warm by 2.4°C (range 1.2 to 4.6°C under IPCC AR6 SSP1-2.6 and SSP5-8.5 scenarios) by 2100 (Good et al. 2019, Kennedy et al. 2019), compared to the average sea surface temperature for 1991–2020. Given that there was no significant temporal trend in late winter SST over our study period, the positive relationship between survival and SST may only hold true under observed conditions. Additionally, extreme events such as winter storms (Frederiksen et al. 2008, Acker et al. 2021) and marine heatwaves (Semmouri et al. 2023), which may increase in frequency with climate change (Cornes et al. 2023), could play an increasingly important role in driving gull population dynamics. Thus, changes in SST during 2002-2017 may have represented a temporary increase in prey resources (e.g., swimming crabs) for gulls but we cannot know the consequences of future increases in ocean temperatures. Thanks to international legislation, background levels of organic pollutants such as organochlorines are predicted to decline, hopefully limiting their negative impacts on top predators like seabirds.

As we found positive effects of ocean temperatures over the study period and as data of pollution and individuals’ body masses were only available from a single year (per individual spanning two years), which may, in part, explain the limited effects of pollution on survival rates, these two stressors cannot explain the pronounced negative trend in survival rates and population sizes observed in this population. This gull population and others along the coast of Norway are undergoing rapid declines and should be of extreme concern in terms of their long-term viability and risk of extinction. The North Sea is also under pressure from other human activities (e.g., commercial fishing, offshore wind farms, oil and gas production and shipping), all known to be important threats to seabirds, particularly during the non-breeding season. Furthermore, here, we focused on core non-breeding areas however, individuals from this population also utilise other areas which are affected by e.g., climate change, fishing competition to differing degrees, which can also impact their survival and population trend. While ocean warming is one potential threat, there are several others which may explain the observed declines in adult survival and breeding population sizes. This emphasises the need for monitoring data that captures changes in anthropogenic pressures and wildlife populations, at a sufficient temporal and spatial resolution, and use such models as used here, to provide predictions of future population viability under ecosystem change.

## Acknowledgements

Funding to conduct the analysis was provided by the Norwegian Institute for Nature Research (SATS Project “CLIMTOX” 18261700) and we wish to acknowledge the project CLEAN, funded by the Fram Centre, Norway. We would like to thank all students and other field assistants for their work in the field over the years, and John A. Henden, Øystein Miland and Øystein Varpe specifically for their assistance during the field campaigns in 2001 and 2002.

## Data Availability Statement

The data that support the findings of this study will be made openly available in Dryad on publication.

