## Supplementary Information S1-3 for "Path analysis reveals combined winter climate and pollution effects on the survival of a marine top predator"

### Contents

Supplementary information S1. Organochlorine (OC) variables and results from principal component analysis ........ 2

Supplementary information S2. Baseline survival model selection ........ 6

**Supplementary information S1.** Organochlorine (OC) variables and results from principal component analysis

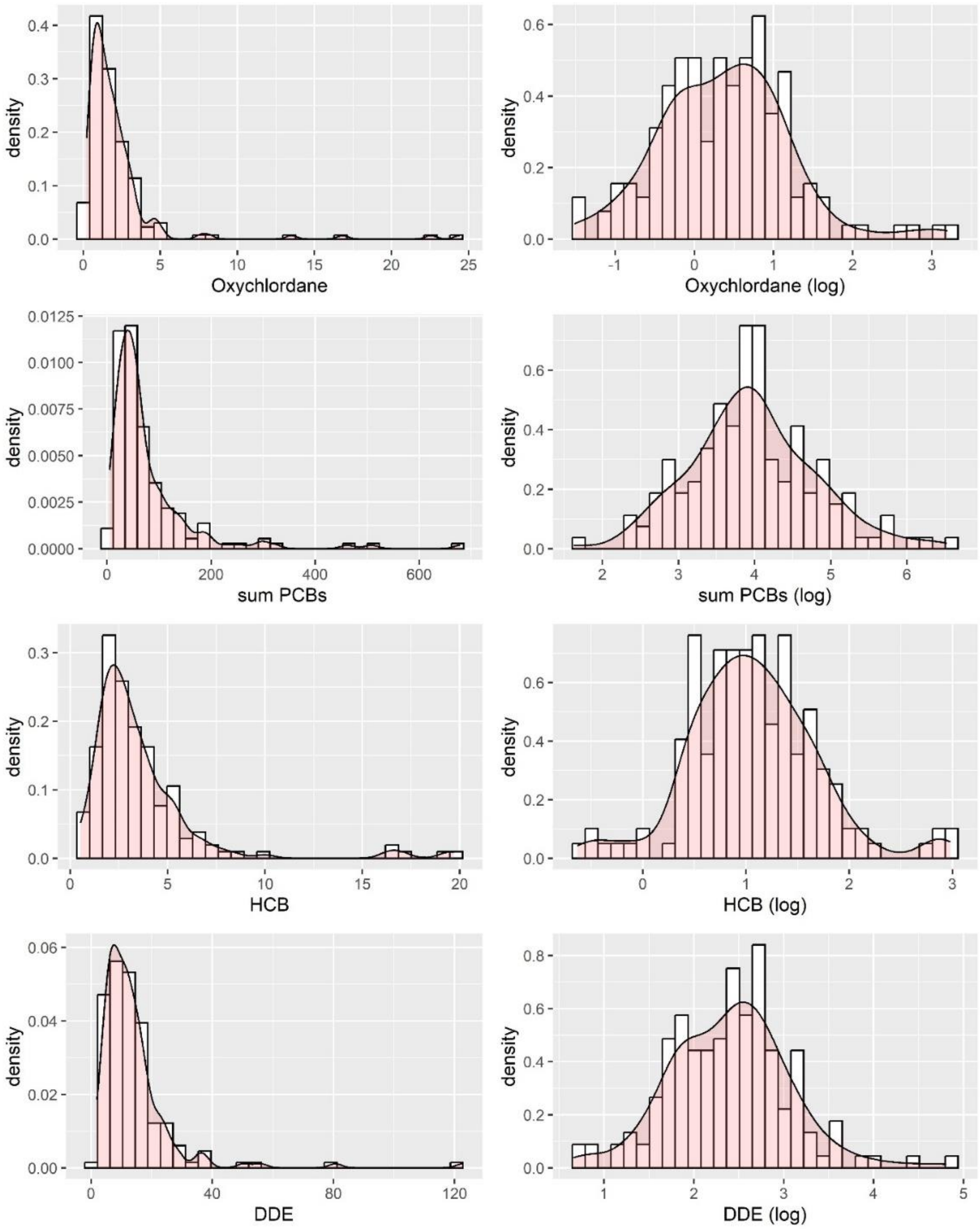


**Figure S1.** Historgrams with density plots of OC levels for the different compounds; Oxychlordane, summed PCBs, HCB and DDe. In the left panel are the measured values of all variables, on the right are the log-transformed data.

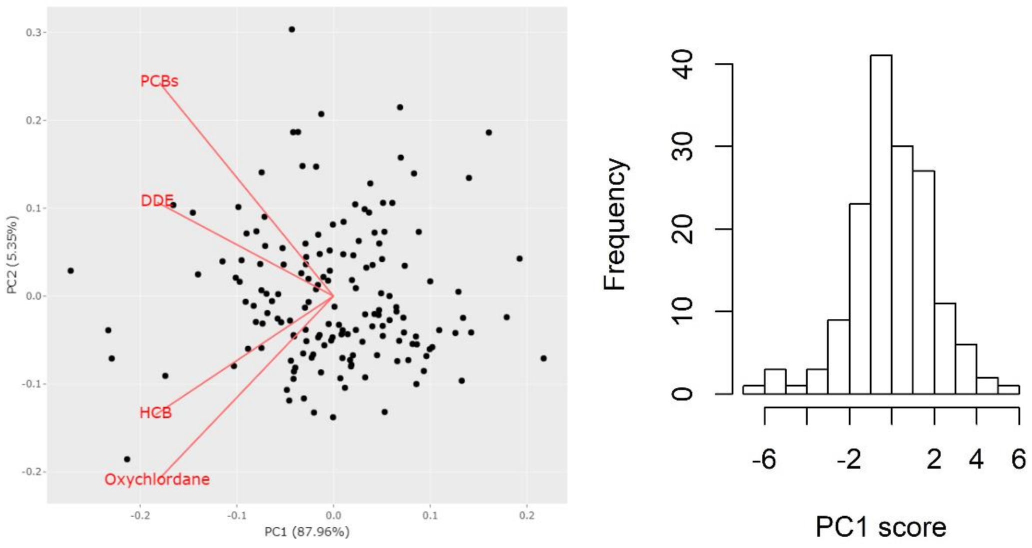


**Figure S2.** (Left) Ordination plot showing the relationship between variables based on a principal component analysis, including individual, log-transformed and scaled variables of summed PCBs, DDE, HCB and oxychlordane blood concentrations, where PC1 explaned 87.96% of the variance. (Right) histogram of the first principal component (PC1) coefficient.

**Supplementary information S2.** Extraction of climate covariates from core foraging areas and seasonal aggregation

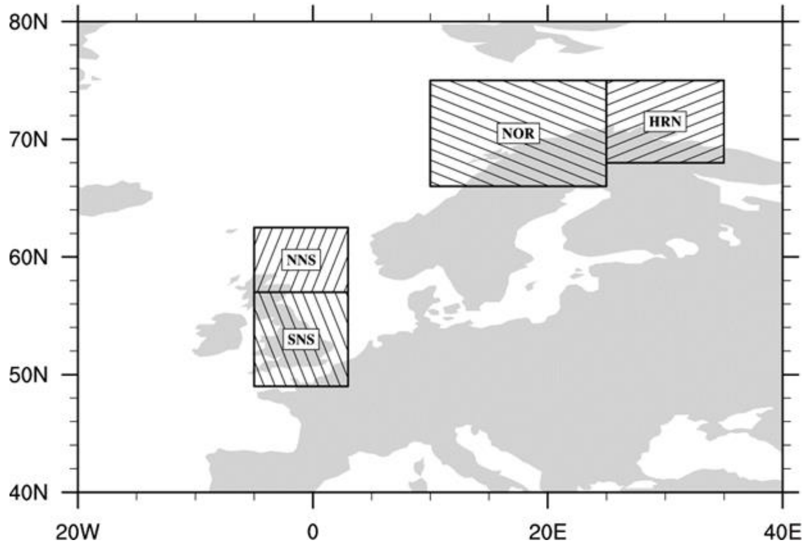


**Figure S3.** Areas from which climate covariates (MSLP, SST and 2m (air) temperature) were extracted, where covariates were extracted from the whole North Sea area (NNS and SNS) during autumn and winter and the area around Hornøya (HRN) during spring.

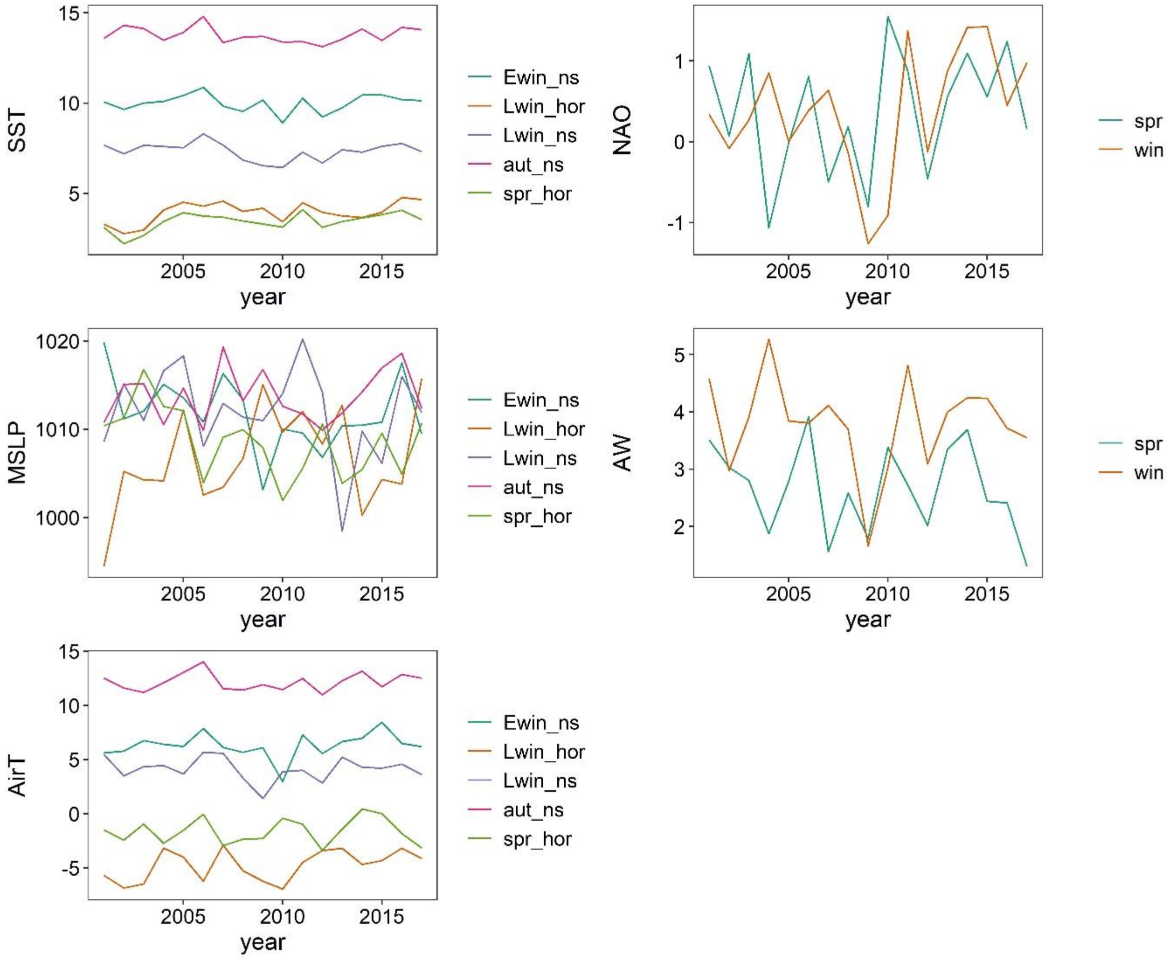


**Figure S4.** Time series of climate covariates. Lines represent values, for the periods used in the analysis, of sea surface temperature (SST), mean sea level pressure (MSLP), air temperatre at 2m (AirT), North Atlantic Oscilliation (NAO) and Atlantic Water inflow (AW).

**Supplementary information S3.** Baseline survival model selection

Models were fitted with a linear year effect (Time), a fixed year effect (time) a trap dependence effect (td, difference in recapture probability between first resighting and later resightings), transience (trans, difference in survival between first resighting year and later resightings) and sex (male/female).

**Table S1.** Model selection table to identify the base model of adult apparent survival rates.trend reflects a temporal trend effect, sex is a male-female contrast, year reflects a fixed year effect and TD refers to a one-year trap dependence effect.

| Survival model | Recapture model | npar | AICc | ΔAICc | weight |
|---|---|---|---|---|---|
| trend | year + TD | 18 | 1154.33 | 0.00 | 0.44 |
| sex + trend | year + TD | 19 | 1154.86 | 0.53 | 0.34 |
| - | year + TD | 18 | 1162.64 | 8.31 | 0.01 |
| sex | year + TD | 19 | 1163.49 | 9.16 | 0.00 |
| sex | year + TD | 19 | 1163.49 | 9.16 | 0.00 |
| trend | year | 16 | 1192.61 | 38.27 | 0.00 |
| sex + trend | year | 17 | 1193.11 | 38.78 | 0.00 |
| - | year | 15 | 1197.06 | 42.73 | 0.00 |
| sex | year | 16 | 1197.86 | 43.53 | 0.00 |
| sex | year | 16 | 1197.86 | 43.53 | 0.00 |
| trend | trend + TD | 5 | 1266.51 | 112.18 | 0.00 |
| trend | td | 4 | 1266.71 | 112.38 | 0.00 |
| sex + trend | trend + TD | 6 | 1267.20 | 112.87 | 0.00 |
| sex + trend | td | 5 | 1267.40 | 113.07 | 0.00 |
| - | trend + TD | 4 | 1267.48 | 113.15 | 0.00 |
| sex | trend + TD | 5 | 1268.29 | 113.95 | 0.00 |
| sex | trend + TD | 5 | 1268.29 | 113.95 | 0.00 |
| - | td | 3 | 1273.25 | 118.92 | 0.00 |
| sex | td | 4 | 1274.23 | 119.90 | 0.00 |
| sex | td | 4 | 1274.23 | 119.90 | 0.00 |
| trend | sex | 4 | 1328.59 | 174.26 | 0.00 |
| sex + trend | sex | 5 | 1329.79 | 175.46 | 0.00 |
| - | sex | 3 | 1333.44 | 179.10 | 0.00 |
| sex | sex | 4 | 1334.79 | 180.46 | 0.00 |
| sex | sex | 4 | 1334.79 | 180.46 | 0.00 |

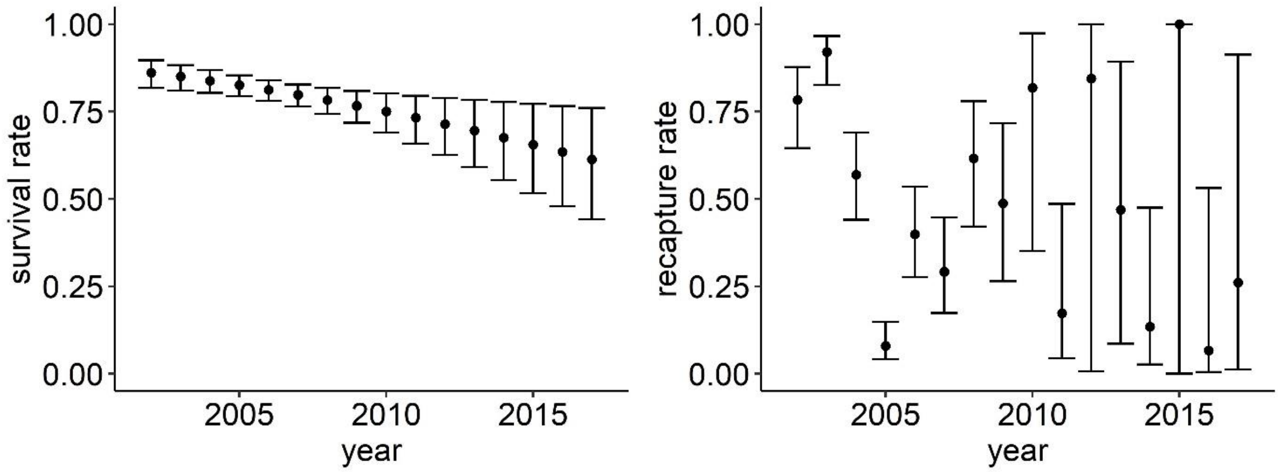


**Figure S5.** Estimates of survival and recapture from best-fitting base model, including a linear trend effect on survival and fixed effects of year (time) and trap dependence (td) on recapture probabilities.

**Table S2.** Candidate models of climate covariate effects on adult survival, based on de-trended (and scaled) covariates. For all models the recapture model was fitted with a time and on-year trap dependence effect. Mean slope estimates are included for the covariate effects. Effects of late winter sea surface temperature in the North Sea ($SST_{Lwin}$) were still statistically significant and positive and the single covariate model was now a better fit that a model also including spring mean sea level pressure around Hornøya colony ($MSLP_{spr}$). All models that had an AICc at least 2 units less than the intercept in the non-detrended analysis were also a better fit in the de-trended analysis. Thus, we confirm that effects are not spurious due to the presence of temporal trends in covariates.

| **Survival model** | **npar** | **AICc** | **ΔAICc** | **weight** | $\mathbf{R^2}$ |
|---|---|---|---|---|---|
| $0.43\mathbf{SST_{Lwin}}$ | 17 | 1153.74 | 0 | 0.38 | 0.27 |
| -0.09**trend** | 18 | 1154.33 | 0.59 | 0.28 | 0.33 |
| $0.37\mathbf{SST_{Lwin}} + 0.20\mathbf{MSLP_{spr}}$ | 18 | 1154.67 | 0.94 | 0.24 | 0.32 |
| $0.23\mathbf{MSLP_{spr}}$ | 17 | 1158.80 | 5.06 | 0.03 | 0.07 |
| - | 18 | 1162.64 | 8.91 | 0.00 | |
